# A Reproducible Neurobiology of Depressive Rumination

**DOI:** 10.1101/365759

**Authors:** D.A. Pisner, J. Shumake, C.G. Beevers, D.M. Schnyer

**Affiliations:** Institute for Mental Health Research (IMHR), 116 Inner Campus Dr. Austin, TX 78705; Department of Psychology, University of Texas at Austin, 108 E Dean Keeton St, Austin, TX 78712

**Keywords:** Multimodal, Microstructure, Resting-State, Triple-Network, Depressive Rumination, Reproducibility, Holism

## Abstract

Depressive Rumination (DR), which involves a repetitive focus on one’s distress, has been linked to alterations in functional connectivity of the ‘triple-network’, consisting of Default-Mode, Salience, and Executive Control networks. A structural basis for these functional alterations that can dually explain DR’s persistence as a stable trait remains unexplored, however. Using diffusion and functional Magnetic Resonance Imaging, we investigated multimodal relationships between DR severity, white-matter microstructure, and resting-state functional connectivity in depressed adults, and then directly replicated our results in a phenotypically-matched, independent sample (total *N* = 78). Among the fully-replicated findings, DR severity was associated with: (a) global microstructure of the right Superior Longitudinal Fasciculus and local microstructure of distributed primary-fiber and crossing-fiber white-matter; (b) an imbalance of functional connectivity segregation and integration of the triple-network; and (c) ‘multi-layer’ associations linking these microstructural and functional connectivity biomarkers to one another. Taken together, the results provide reproducible evidence for a multi-layer, microstructural-functional network model of rumination in the depressed brain.

---

Cognitive models of depression posit that negatively biased self-referential processing plays a critical role in maintaining the disorder (Beevers, 2005). Depressive Rumination (DR) is the perseverative form of this cognitive bias, which involves a repetitive focus on “one’s depressive symptoms and on the implications of those symptoms” (Nolen-Hoeksema & Morrow, 1991). DR has been shown to significantly contribute to depressive symptom severity, duration, and relapse likelihood across multiple forms of depression (Nolen-Hoeksema et al., 2008; Nolen-hoeksema, 2000). Despite its central role in depression maintenance, there are few effective treatments for DR (Watkins, 2015; Wells & Papageorgiou, 2008). To gain a precise understanding of underlying vulnerabilities and identify reliable treatment targets, we seek to develop a more unified and reproducible model of DR’s neurobiology using multimodal analysis with replication (Woody & Gibb, 2015).

## A Triple-Network Functional Connectivity Model of Depressive Rumination

At the core of DR is *self-referential processing* (Nejad et al., 2013), which is inwardly directed attention focused on the self. This mechanism is associated with functional activation of modular cortical areas, such as the Precuneus (Yoshimura et al., 2009) along with the anterior and posterior cingulate cortices (ACC; PCC) (Brewer et al., 2013); each of these areas has been associated with self-focused cognitive processes that include thought-suppression, mentalization, and prospection (Buckner & Carroll, 2007; Kühn et al., 2012; Laurita et al., 2017; Ed Watkins & Teasdale, 2001). Although few studies have explored grey matter structural biomarkers of self-reference, reduced cortical volume of the Precuneus is known to be associated with autobiographical memory disturbances (Ahmed et al., 2018; Freton et al., 2014) and poor metacognition (McCurdy et al., 2013; Ye et al., 2018). Since self-referential processing is believed to predominantly operate in a task-negative brain state (Rosenbaum et al., 2017), moreover, studies that employ resting-state functional MRI (rsfMRI) uniquely offer some insight into the neurobiology of self-reference (Nejad et al., 2013; Northoff et al., 2006). Indeed, rsfMRI has enabled researchers to more broadly attribute self-referential processes to the Default-Mode Network (DMN), whose functional connectivity may reflect mind-wandering (Greicius et al., 2003; Rosenbaum et al., 2017) and for which the Precuneus is a core hub (Utevsky et al., 2014). Precise features that differentiate “healthy” mind-wandering from maladaptive ruminative patterns are difficult to distinguish, however, perhaps due to the inherent subjectivity of inner mentation along with fundamental limitations of the resting-state modality in general (Whitfield-Gabrieli et al., 2011). Nevertheless, increased DMN activity during internally-focused attention is generally believed to amplify self-referential processing and thereby increase the demand on cognitive resources (Leech et al., 2011; Sheline et al., 2009). In particular, recent studies have shown that greater DR severity is associated with lower functional connectivity within a parietal sub-system of the DMN, the Precuneal DMN (pDMN), which is both vulnerable to malfunctioning under conditions of cognitive resource depletion (Wang et al., 2016), and specifically responsible for regulating the episodic memory retrieval that supports inner mentation (Rosenbaum et al., 2017).

*Emotional salience*, a second proposed mechanism of DR, may be necessary for assigning a negative bias to self-referential processing. A wealth of literature has shown that emotional salience is associated with both the function and structure of the ACC, Insula, and Amygdala, and can extend to other regions as well (Jilka et al., 2014; Liang et al., 2015). Greater sustained activation within the Amygdala has been observed, for example, in response to negative-valence words following non-emotional words in depressed individuals (Siegle et al., 2002). Whereas the amygdala may act as a source of emotional signaling (LeDoux, 2003), the ACC assigns salience to emotional signals in the service of goal attainment (Etkin et al., 2011). In the context of DR such goals might include relief from depressive symptoms, for instance. Research into the precise role of the ACC in DR have been mixed, however (Cooney et al., 2010; Kühn et al., 2012; Nejad et al., 2013). On the one hand, discrepancies across studies may reflect the variety of motivational states that are engaged during different task paradigms (Kolling et al., 2016). On the other hand, they might also allude to its dense interconnectivity with other cortical areas (Etkin et al., 2011) or the multiple specialized functions among sub-regions of the ACC itself (Stevens et al., 2011). Thus, emotional salience in DR may be more precisely understood within a functional connectivity perspective of the Salience Network (SN)(Ordaz et al., 2016). In fact, a recent study on depressed teenagers found a critical role for the SN in the maintenance of negative cognitive bias and self-criticism in DR^57^. One reason for this finding may be that aberrant functional connectivity of the SN during self-referential processing results in misappropriated “tagging” of self-relevant thoughts with a negative bias (Nejad et al., 2013), perhaps through dysregulated reward circuitry (Yang et al., 2016). Along these lines, a recent task-based fMRI study showed that depressed ruminators tend to more actively recruit SN nodes simultaneously during reward anticipation. Hence, emotional salience in the context of DR might also or alternatively be construed as a hypervigilance to self-relevant cues (Kocsel et al., 2017).

A third proposed mechanism of DR is impaired *attentional engagement*, which may instantiate repeated failures to disengage from negatively-salient self-referential processing while also preventing attentional shifts toward neutral or positive thoughts (Southworth et al., 2017). In depressed individuals, this impaired disengagement (Grafton et al., 2016; Koster et al., 2011; Southworth et al., 2017) is believed to amplify and sustain the negative bias of self-referential processing by spurring a vicious cycle of self-criticism that drains the cognitive resources needed to counteract the increase (Bernstein et al., 2017). Thus, the mechanism of impaired disengagement may lend explanation to the core perseverative feature of DR, which distinguishes it from more adaptive, reflective forms of rumination (Nolen-Hoeksema et al., 2008; Rude et al., 2007). Broadly construed, impaired disengagement in DR likely implicates the aberrant structural and functional properties of cognitive control modules, such as the dorsolateral and inferior prefrontal cortices (Jacobs et al., 2016; Kühn et al., 2012; Wang et al., 2015), whose activation patterns are largely titrated by the ACC (Etkin et al., 2011). The functional connectivity perspective of the Executive Control Network (ECN), which considers these frontal modules as a collection of network subunits, may therefore more parsimoniously capture the broader role of impaired disengagement in DR.

Perhaps in light of the three proposed mechanisms of DR, the three corresponding resting-state networks (RSNs) previously discussed have become ubiquitous in the study of DR’s functional neurobiology—the DMN, SN, and ECN (Bernstein et al., 2017; Ordaz et al., 2016). Nevertheless, these RSN’s have been studied largely in isolation – i.e. with respect to their *within-network* functional connectivity patterns –in the context of DR research (Nejad et al., 2013; Ordaz et al., 2016). Despite their diverse specialized intrinsic functions, however, mounting research has shown that the DMN, SN, and ECN also interact extrinsically with one another (Menon, 2011). Hence, additional insights might be gleaned by studying these particular RSN’s as a mesoscale system of networks – the “triple network” – as some have already proposed more broadly in the study of depression (Liu et al., 2017; Menon, 2011; Wu et al., 2016; Zheng et al., 2015) (*See* **Figure 1**). In fact, only one study to date has explored extrinsic, *between-network* functional connectivity patterns in DR. Specifically, Hamilton et al. (2011) used state-change analysis of rsfMRI to show that higher DR severity is associated with greater DMN dominance over the ECN, with a variable role for the SN across differing levels of depression severity. That study importantly did not include structural brain measures, however, which are still needed to explain DR’s persistence as a stable trait (Brosschot et al., 2006).

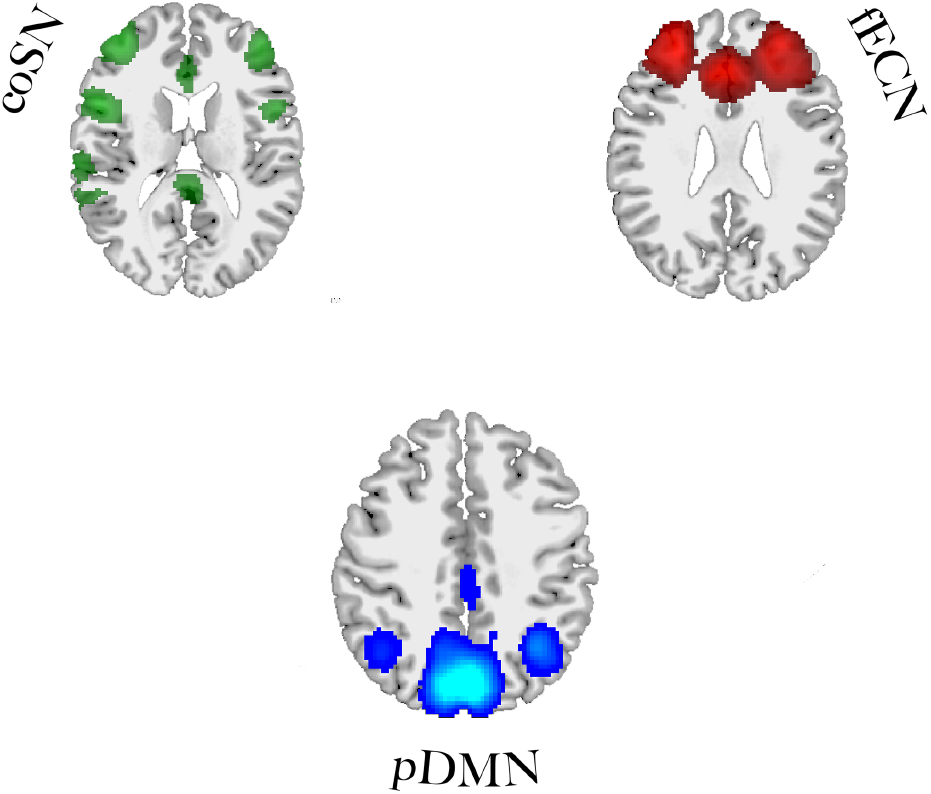

## A Microstructural Model of Depressive Rumination

Studies have shown that inter-individual differences in DR severity likely indicate the existence of structural brain correlates of DR (Kühn et al., 2012; Nejad et al., 2013; K. Wang et al., 2015; Zuo et al., 2012). As a basis for this, the perseveration implicated in DR is believed to be the neurodevelopmental expression of a stable trait, ingrained across brain structures over time (Brosschot et al., 2006). Hence, it would also seem to implicate a complex structural basis, but one that, like in the functional case, cannot be explained modularly through grey matter cortices alone (Wang et al., 2015). It follows that the structural analogue to functional connectivity – microstructural connectivity (Peer et al., 2017) – may be a useful framework for studying the structural basis of trait DR in particular. Microstructural connectivity is typically studied using methods like diffusion tensor MRI (dMRI), which provides an *in vivo* method for measuring global and local Fractional Anisotropy (FA) based on anisotropic properties of WM in the brain (Yendiki et al., 2011). In fact, BOLD signal clustering of resting-state WM has been shown to generate patterns resembling white-matter tracts (Mezer et al., 2009). Although recent studies beyond DR literature have shown that WM microstructure of frontoparietal tracts, such as the Cingulum (CCG) and the Uncinate Fasciculus (UF), may support the DMN (Tao et al., 2015) and ECN (Steffens et al., 2011) respectively, only one study to date has explicitly investigated the WM basis for DR. Namely, Zuo et al. (2012) used Tract-Based Spatial Statistics (TBSS) to show that FA of the Superior Longitudinal Fasciculus (SLF) and neighboring motor fibers is negatively associated with DR. Due to its small sample size (n = 16), however, those results may not generalize to other depressed populations. Still, multimodal attempts have not been made to reconcile these or other WM biomarkers with the more abundant functional biomarkers identified from fMRI literature.

## A Microstructural-Functional Model of Depressive Rumination

To review, DR’s neurobiology is likely characterized by a variety of neural biomarkers that operate across multiple levels of neural complexity, correspond to at least the three proposed cognitive mechanisms of DR, and encompass both functional and structural dimensions of the brain. Although older studies of modular biomarkers have identified a core set of nodes relevant to each of these mechanisms, the modular perspective cannot account for dynamic interactions across those nodes (Monnart et al., 2016). More recent unimodal connectivity research has also failed to address whether the tentative set of functional and structural connectivity biomarkers of DR already identified represent independent or interdependent (Menon et al., 2011) brain systems. Hence, a more ‘holistic’ exploration of ‘multilayer’ connectivity biomarkers (Braun et al., 2018), identified across both functional (Hamilton et al., 2015; Ordaz et al., 2016) and structural (Zuo et al., 2012) modalities (Chavez & Heatherton, 2013; Medaglia et al., 2016), may be maximally informative for building more precise models of rumination in depression. Likewise, neuroscientific appeals made to either structural or functional connectivity biomarkers of DR, considered in isolation, might reduce the precision of our etiological understanding (Mazzocchi, 2012). To therefore clarify any systems-level (Bassett & Gazzaniga, 2011) mechanisms responsible for maintaining DR as a stable trait (Bagby et al., 2004; Fawcett et al., 2015; Mandell et al., 2014), multimodal studies of functional and microstructural connectivity, considered in tandem, are needed.

In the present study, we propose two hypotheses: 1) Microstructural connectivity biomarkers and triple-network functional connectivity biomarkers of depressive rumination (DR) exist, where the latter corresponds to each of three proposed cognitive mechanisms of DR—aberrant self-referential processing, negatively-biased salience processing, and impaired disengagement of attention; 2) inter-individual differences in trait DR *severity* can be explained through joint appeal to triple-network functional connectivity with respect to underlying microstructural connectivity constraints. To test and explore these hypotheses, we utilize data from two neuroimaging modalities, dMRI and rsfMRI. Given the absolute importance of replication in cognitive neuroscience (Button et al., 2013) and the growing need for more credible and reproducible neurobiological models of disordered cognition (Mulugeta et al., 2018), we additionally perform our analyses across two separate datasets, acquired independently and at different locations. This approach, which is rarely used in social or biological sciences (Button et al., 2013; Poldrack et al., 2017), would allow us to determine whether initial discovery findings could be directly replicated in an independent sample. The findings that did replicate would, in turn, afford a higher degree of confidence in support of our conclusions, and perhaps provide an auxiliary means of inferring generalizability beyond statistical power alone.

We test our first hypothesis by applying multiple linear regression analyses to dMRI and rsfMRI data independently. The dMRI analyses include both TBSS and global probabilistic tractography methods (Yendiki et al., 2011), whereas the rsfMRI analyses encompass both within-network and between-network (Parlatini et al., 2017) resting-state functional connectivity based on Independent Components Analysis (ICA)(Ordaz et al., 2016). We then test the second hypothesis by examining joint microstructural-functional connectivity relationships associated with DR. Lastly, we test both of our hypotheses by applying a near-identical analytic pipeline developed for the discovery sample to an independent sample obtained from NKI Rockland (Nooner et al., 2012, *RRID:SCR_009435*).

## METHODS

### Participants – Discovery sample

Thirty-nine treatment-seeking participants with DSM-IV Major Depressive Disorder (MDD) were recruited for this study from advertisements placed online, in newspapers, and on late night TV. Participants were screened for medical or physical conditions that would preclude participation in an MRI study. They also completed an abbreviated Mini International Neuropsychiatric Interview (MINI)(Sheehan et al., 1998) to determine provisional MDD diagnosis, which were then confirmed using in-person Structured Clinical Interviews for the DSM-IV Disorders (SCID)(First et al., 1997) administered by a trained research assistant. Participants were excluded if they met criteria for past year substance abuse or dependence, current or past psychotic disorder, bipolar disorder, and schizophrenia. Participants receiving pharmacological treatment were allowed into the study if there has been no medication change in the 12 weeks prior to study entry. To minimize brain changes associated with aging, participants were between ages 18-55 (Beevers et al., 2015).

### Ethics Statement

The Institutional Review Board at the University of Texas at Austin approved all study procedures and materials and all participants provided signed informed consent.

### Depressive Rumination Measurement

The RSQ (Response Styles Questionnaire)(Erdur-Bakera & Bugaya, 2010) is a 10-item self-report measure of the tendency to ruminate. It consists of a total score and two sub-scales: reflection and brooding. The reflection subscale measures an individual’s tendency to turn inward to engage in problem-solving and thereby alleviate negative or depressed mood (Nolen-Hoeksema et al., 2008), whereas brooding measures the maladaptive form of rumination believed to be a proxy for DR (Hamilton et al., 2011; Nolen-Hoeksema et al., 2008). Brooding specifically reflects the intensity of ruminative responses to expressions of negative emotion (Nolen-Hoeksema & Morrow, 1991). For each item, subjects indicate the frequency of each event on a scale ranging from 0 (“almost never”) to 3 (“almost always”), yielding a range of possible scores from 0-30. The brooding subscale has high reliability (a=0.77-0.92)(Erdur-Bakera & Bugaya, 2010), is well-validated within depressed populations (Treynor et al., 2003), decontaminated of any explicitly depressive content (Treynor et al., 2003), and the sub-scale of choice for most neurobiological studies of DR (Hamilton et al., 2011; Nejad et al., 2013; Ordaz et al., 2016; Rosenbaum et al., 2017; Southworth et al., 2017). For these reasons, we used the brooding subscale exclusively to measure DR.

### Depression Severity Measurement

The Beck Depression Inventory (BDI)(Beck et al., 1996) is a 21-item self-reporting questionnaire for evaluating the severity of depression in normal and psychiatric populations. It contains 21 items on a 4-point scale from 0 (symptom absent) to 3 (severe symptoms), and instructs the participant to recall depression symptoms occurring over the previous two weeks.

### Imaging Acquisition

MRI scans were acquired on a whole body 3T GE MRI with an 8-channel phase array head coil. The scanning protocol involved collection of a localizer followed by a high-resolution T1 structural scan, two resting state scans of 6 minutes each, a second high-resolution structural scan, and finally a 55-direction diffusion tensor (dMRI) scan. For the resting-state scan, instructions were presented utilizing a back-projection screen located in the MR bore and viewed through a mirror mounted on the top of the head coil. Participants were instructed to remain awake and alert and keep their gaze on a fixation cross (+) presented approximately at the center of their field of view for the 6-minute duration of the scan. (*See* **Appendix, Methods: Section A**).

### dMRI Preprocessing

Preprocessing of Diffusion Magnetic Resonance Imaging (dMRI) data was carried out using a custom preprocessing workflow that included eddy correction, brain extraction, denoising, and tensor/ball-and-stick model fitting tools adapted from the FMRIB Diffusion Toolbox (Jenkinson et al., 2012, *RRID:SCR_002823*, http://www.fmrib.ox.ac.uk/fsl/). To achieve maximal sensitivity and specificity from the dMRI data, preprocessing included rigorous automated and manual quality control steps (*see* **Appendix, Methods: B)**.

### dMRI: Tract-Based Spatial Statistics (TBSS) / Tractography and DR

A number of approaches to dMRI analysis were implemented. To begin, the whole-brain data was interrogated using Tract-Based Spatial Statistics (TBSS)(Smith et al., 2006) to identify microstructural characteristics that were associated with brooding severity (*see* **Appendix, Methods: Section C**). We additionally employed the ‘crossing-fibers’ extension of TBSS (Jbabdi et al., 2010), which is less often used due to its computational expense, but can uniquely capture group differences in secondary (i.e. “crossing”) fibers, unlike TBSS based on the tensor model alone.

For statistical testing, a permutation approach was employed using FSL’s “randomise” function with the TFCE Threshold-Free Cluster Enhancement option, generating 10,000 permutations and applying family-wise error (FWE)-correction to obtain cluster inferences. A two-tailed regression model was next generated using FSL’s GLM function, whereby RSQ brooding scores were used as the criterion variable with age and gender as nuisance covariates. Age was included due to a well-known confounding influence of age on WM microstructure (Westlye et al., 2010), and gender was included due to some evidence of gender differences in rumination—namely, that females tend to exhibit greater rumination severity than males (Nolen-Hoeksema et al., 1987). The resulting negative contrast mask (i.e. representing those voxels negatively associated with greater DR severity) was thresholded twice—first at *p*=0.05 and again at *p*=0.01. The use of two thresholds allowed us to determine those voxels which were most strongly associated with DR (*p*=0.01), while also yielding a more liberal cluster of voxels (*p*=0.05) that could be reliably used as a Region-of-Interest (ROI) for multimodal analysis.

Following TBSS, we sought to corroborate our initial group level, voxel-wise dMRI findings using individual-level tractography, which is an alternative dMRI methodology that attempts to reconstruct known WM pathways while retaining each subject’s image in native space orientation. Since spatial information is not manipulated in tractography as it is with TBSS (De Groot et al., 2013), tractography could confirm any TBSS findings in native space, while also disconfirm false-positives due to non-physiological factors such as image misalignment, movement, and other factors resulting from the methodological limitations of TBSS (Torgerson et al., 2013).

For tractography, we chose to analyze average FA measures from the entire pathways of tracts of interest whose labels included significant voxels from the earlier TBSS stage. These measures were then further analyzed across hemispheres to establish any significant laterality effects (Vernooij et al., 2007). To perform tractography, we specifically used the TRActs Constrained by UnderLying Anatomy (TRACULA, *RRID:SCR_013152*, http://surfer.nmr.mgh.harvard.edu/fswiki/Tracula) tool in FreeSurfer (version 5.3.0, *RRID:SCR_001847*, http://surfer.nmr.mgh.harvard.edu/) (Yendiki et al., 2011), which delineates 18 known WM bundles using each participant’s joint dMRI and T1-weighted MRI data (*See* **Appendix, Methods: Section D**).

### rsfMRI Preprocessing

Preprocessing of baseline rsfMRI data was carried out using a custom workflow based on FSL’s FEAT (Jenkinson et al., 2012), combined with AFNI and FREESURFER tools. Additional denoising steps were taken by controlling for WM and ventricular CSF confounds, and by using FSL’s ICA-based Xnoisifier artifact removal tool (FIX) to control for motion and physiological artifact (*See* Appendix, Methods: **Section E**).

### rsfMRI: Group Independent-Components Analysis (ICA) / Dual-regression and DR

Group-level Independent Components Analysis (ICA) was performed by employing “temporal concatenation” of the complete, preprocessed rsfMRI time-series from all of the participants and restricted to twenty-five independent component (IC) outputs (Beckmann et al., 2009). Four IC’s were manually identified as noise and removed from further examination. Of the remaining twenty-one networks, all were identified using visual inspection by way of reference to the 17 RSN’s delineated by the Yeo et al. 2011 atlas (Yeo et al., 2011), thus allowing for identification of the three IC’s of the triple-network that were introduced above - the pDMN (Hamilton et al., 2011; Rosenbaum et al., 2017), the coSN (Wu et al., 2016), and the fECN (Bernstein et al., 2017; Dutta et al., 2014; Watkins & Brown, 2002).

A dual-regression approach (Smith et al., 2012) was next performed on the triple-network RSN’s, which were used as regressors for each individual subject’s 4D set of fMRI volumes in order to extract time-series that were both specific to each subject and to each of the three IC’s^64^. Design matrices and contrasts were then created to test for correlations between brooding severity and total average intrinsic connectivity within each of these RSN’s, controlling for age and gender. These regression models were tested separately for each RSN, using two-tailed contrasts in an identical manner to that used in TBSS, with FSL’s randomise (10,000 permutations) and TFCE cluster-thresholding with whole-brain FWE-correction (Smith & Nichols, 2009). To further correct for analysis-level multiple comparisons among the three triple-network RSN’s, we also Bonferroni-corrected our alpha significance level for these models to 0.05/3 or *p*=0.0167.

### Between-Network Functional Connectivity and DR

To investigate interactions between the triple-network RSN’s, we utilized FSLNets (Smith et al. 2013, https://fsl.fmrib.ox.ac.uk/fsl/fslwiki/FSLNets), a MATLAB-based tool that interfaces with FSL. FSLNets utilizes the individual resting-state network IC’s generated from the earlier dual-regression stage to perform hierarchical clustering across RSN’s and estimate a partial-correlation (Spearman’s *ρ*) matrix for each RSN for each participant. To perform between-RSN general linear modeling, randomise was again employed with FWE-correction, but this time using the nets_glm function at 10,000 permutations. The design matrices and contrasts used in the earlier FSL GLM analyses, were used in this analysis as well. In summary, whereas the dual-regression approach identified DR-associated clusters (both *belonging to* and *external to* the RSN’s) that are more strongly connected to each of the triple-network RSN’s considered separately, the FSLnets approach identified DR-associated patterns of between-network correlation among the triple-network RSN’s considered collectively.

### Replication and Statistical Power

To formally test whether the findings would generalize beyond the discovery sample (Poldrack et al., 2017), we directly replicated all stages of analysis using a phenotypically similar, independent sample obtained from the publicly-available, multi-site NKI Rockland dataset (Nooner et al., 2012, *RRID:SCR_009435*). By chance, that dataset contained a sub-sample of participants with equivalent demographics to those participants from our original sample; this included (N=39) participants who were between the ages of 18-55, had no history of drug abuse or severe comorbid psychopathology, had both useable dMRI data and rsfMRI data, and reported at least some depressive symptomatology. The participants had also been administered the 21-item Beck Depression Inventory (BDI-II) and the 22-item Rumination Response Scale (RRS)(See **METHODS: Depressive Rumination Measurement**), which contained identical brooding subscale items to those administered in the 10-item RSQ scale (Treynor et al., 2003). As part of a larger battery of measures, many but not all participants also completed the Structured Clinical Interview for DSM-IV-TR Axis I Disorders (SCID)(First et al., 1997). Specifically, 33% of the n=39 participants met full criteria for MDD (18% current/recurrent and 15% in remission), requiring that the remaining participants be included on the basis of broader depressive symptomology. This included 66% with dysphoria as determined by elevated BDI-II scores (≥4) — a cutoff used in prior studies (Berle & Moulds, 2013; Williams & Moulds, 2007). One advantage to including dysphoric participants was that it could possibly allow us to infer whether any replicated biomarkers of DR are specific to DR and not merely conflated with the depressive symptoms specific to an MDD diagnosis.

Furthermore, to minimize the possibility of sample-variant effects due to incongruence of neuroimaging acquisition parameters across samples (Gouttard et al., 2008), we spatially and temporally resampled the rsfMRI and dMRI data in the replication sample to match the voxel resolution of the rsfMRI and dMRI data and the sampling frequency (i.e. TR) of the rsfMRI data in the discovery sample (*See* **Appendix, Methods: Sections F & G**). To ensure direct equivalence of neuroimaging data preprocessing in the replication (Bowring et al., 2018), we also applied a nearly equivalent analytic pipeline to that applied to the discovery sample, but using the pDMN, fECN, and coSN group-ICA RSN definitions from the discovery dataset. Lastly, given the multiple scanner sites used to collect the neuroimaging data in the Rockland sample, we employed mixed-effects regression models (both with FSL’s GLM and in R 3.4.0, *RRID:SCR_001905*), whereby scanner site was additionally modeled as a random effect.

Since generalizability was a core aim of our study, power analyses were also conducted for detecting a large effect size when using multiple regression with two covariates (age and gender) and a significance level of α<0.05 (*See* **Appendix: Methods, Section H**). Given our sub-optimal sample-sizes, we also opted to filter significant findings by a large effect size cutoff of R^2^=0.25 for non-voxel-wise tests (i.e. tractography, multimodal analysis of beta-coefficients) and *p*<0.01 FWE-corrected threshold for voxel-wise tests (TBSS, Dual-Regression, FSLnets). This step would serve to more stringently identify those findings with the highest putative generalizability as well as those at risk of replication failure. Further, we classified a finding as being a ‘full replication’ if it replicated with respect to both equivalence in directionality of the effect and the RSN’s or anatomical location(s) implicated. Likewise, the fully-replicated findings needed to meet our effect-size cutoff in at least one of the two samples, as well as an FWE-corrected α<0.05 level of significance or uncorrected α<0.001 level of significance for ROI analyses. Finally, to evaluate analysis-level false-discovery across multimodal replications, we further generated a null-distribution for meta-analysis based on replication counts (*See* **Appendix: Methods, Section I**).

## RESULTS: INITIAL SAMPLE

### Descriptive Statistics and Behavioral Findings

Depressive rumination severity (DR) as measured using the RSQ was normally distributed (*M*= 9.08, *SD*= 3.01, range: 17-48, IQR=5; Shapiro-Wilk = 0.976, *p* = 0.53). Depression severity was also normally distributed (*M*= 31.94, *SD*= 8.16, range: 1-15, IQR=10, Shapiro-Wilk = 0.976, *p* = 0.35) and was modestly associated with DR severity (R^2^ = 0.13, *F*(1, 37) = 5.61, *p*=0.02). Age was positively skewed (*M*=27.51, *SD* = 8.80, range: 18-52) due to an overrepresentation of young adults in our discovery sample. Although age was weakly correlated with RSQ brooding scores (R^2^ = 0.09, *F*(1, 37) = 3.84, *p*<0.06) and interacted with several neuroimaging measures throughout our analyses, no significant interactions with gender were observed in either discovery or replication samples. Nevertheless, we retained both age and gender as nuisance covariates for each regression model to ensure maximal generalizability of our findings and consistency in our analyses.

### White Matter (WM) Microstructure and Depressive Rumination

#### TBSS: Localized White-Matter Biomarkers of DR

Our first set of analyses employed tensor and crossing-fiber variations of TBSS to test for WM microstructural connectivity biomarkers of DR (*see* **METHODS: TBSS; Appendix, Methods: Section C**). Results first revealed significantly lower FA in association with higher brooding scores within the right Superior Longitudinal Fasciculus (SLF, both parietal and particularly temporal parts), the Cingulum, and the Corticospinal Tract (CST) (*p*<0.01 FWE; *p*<0.05 FWE; *see blue FA and green F1 clusters in* **Figure 2**). Some of these voxels also extended into the Anterior Thalamic Radiation (ATR), Uncinate Fasciculus (UF), and Splenium (*p*<0.05 FWE; *p*<0.05 FWE; *p*<0.05 FWE). A subsequent ‘crossing-fibers’ TBSS analysis closely tracked these findings, but further revealed significant relationships between DR and the microstructure of secondary-fibers (captured by the F2 partial volume) in a small cluster of the right superior Corona Radiata where the SLFT intersects with the Corpus Callosum at the Centrum Semiovale (*p*<0.01 FWE; *see red cluster in* **Figure 2**).

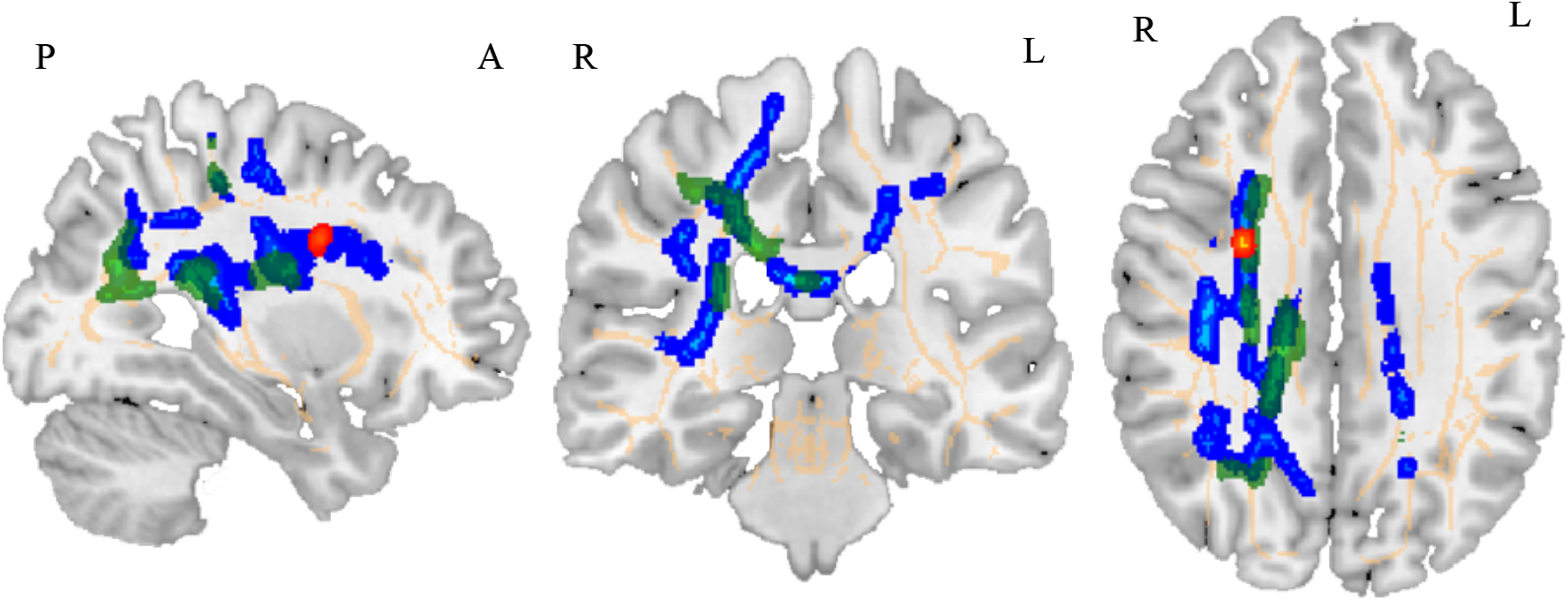

#### Tractography: Globalized White-Matter Biomarkers of DR

Using global probabilistic tractography, we next sought to both methodologically triangulate our TBSS findings and clarify the mereology of DR-associated WM pathways (i.e. their globality vs. locality). Given that TBSS analysis is performed on images where the WM tracks are aligned with an MNI standard, we performed probabilistic tractography on an individual subject basis to provide converging evidence that our findings are not driven by spatial normalization(Yendiki et al., 2011). Using six pathways of interest from the outputs of TRACULA, including the right SLFT (Temporal part), right SLFP (Parietal part), CCG, CST, ATR, and Splenium, and Bonferroni Correction for the six tracts tested (*p*_corrected_=0.05/6 = 0.0083), we found a robust negative association between weighted average FA of the right SLFT and DR severity (adj. R^2^ = 0.18, *F*(3, 36) = 3.85, *p*<0.005). This clarified the findings conducted on a voxel-wise (i.e. spatially-normalized) basis with TBSS, by suggesting that the right SLF may be globally associated with DR severity, whereas the other DR-associated pathways may be better construed as localized WM clusters.

#### Tract Hemisphericity

The aforementioned TBSS and tractography analyses indicated that the SLFT WM finding was distinctly right-lateralized. To test this formally, we treated hemisphere as a within-subjects measure and used Analysis of Variance (ANOVA) to compare two GLM’s predicting DR for each tractography measure of global average FA. Specifically, the first GLM used left hemisphere as the predictor, whereas the second GLM used right hemisphere as the predictor, controlling for left hemisphere. From this analysis, we found supportive evidence for right lateralization of the SLFT (F(36,1)=8.71, *p*=0.005).

### ‘Within-Network’ Functional Connectivity and Depressive Rumination

#### Dual-Regression Findings

We next tested whether DR was associated with resting-state functional connectivity of the three triple-network RSN’s as identified from the outputs of group-ICA followed by dual-regression (*see* **Figure 3** & **METHODS: rsfMRI Group-ICA** & **Dual-regression**). Results revealed that DR was highly correlated with functional connectivity in multiple clusters, some of which belonged to each respective triple-network RSN (i.e. ‘intrinsic’ connectivity), but also others that were outside of the respective RSN’s (i.e. ‘extrinsic’ connectivity) (*see* **Figure 4**). With respect to the latter, we found that DR was positively associated with extrinsic connectivity between the frontal Executive Control Network (fECN) and a small cluster (voxels=15) located at a juncture between the left Amygdala and Parahippocampal Gyrus (*p*<0.05 FWE; *see red clusters in* **Figure 4**). Additionally, the Precuneal Default Mode Network (pDMN) exhibited lower extrinsic connectivity with the left somatosensory areas, left Precentral Gyrus and left inferior Frontal Gyrus (*p*<0.01 FWE; *see blue clusters in* **Figure 3**), but also lower intrinsic functional connectivity between the Precuneus and PCC. Lastly, the Cingulo-Opercular Salience Network (coSN) exhibited lower intrinsic functional connectivity among the dorsal left Precentral Gyrus, left inferior Frontal Gyrus, right superior Postcentral Gyrus, and most prominently in Broca’s area bilaterally (*p*<0.01 FWE; *see green clusters in* **Figure 4**).

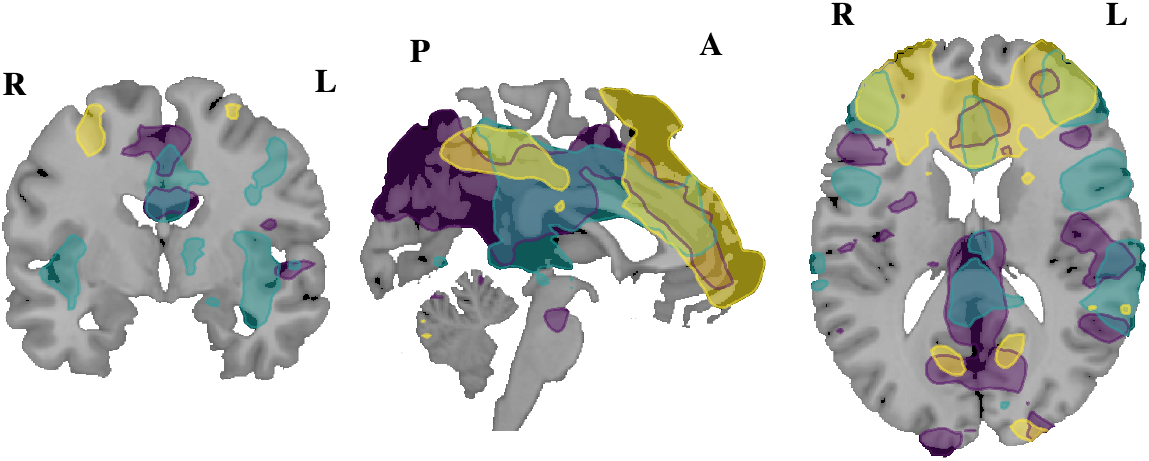

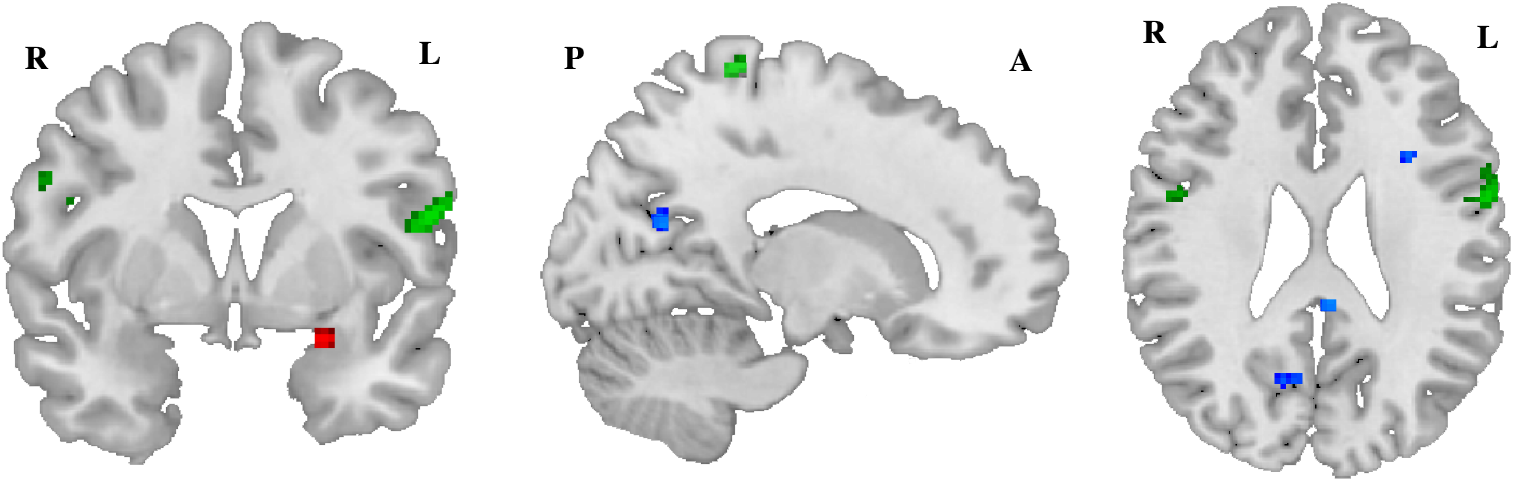

#### WM Microstructural and ‘Within-Network’ Functional Connectivity

We next extracted the beta coefficients representing average total connectivity (intrinsic and extrinsic) for each of the DR-associated triple-network clusters discovered through dual-regression. This step would in turn allow us to test for any multimodal relationships between the within-network functional connectivity findings and the microstructural connectivity findings (i.e. informed by tractography and TBSS). To accomplish this, we first created a matched subsample of n=32 subjects, since seven subjects did not have both useable rsfMRI and TRACULA data. As an initial step, we tested for any associations between the dual-regression clusters and average FA extracted from the thresholded (*p*=0.05) negative contrast mask from TBSS. That analysis revealed that average FA negatively predicted within-network functional connectivity of the pDMN and coSN (adj. R^2^ = 0.14, *F*(3, 29) = 2.75, *p*<0.05; adj. R^2^ = 0.03, *F*(3, 29) = 1.33, *p*=0.06), but not the fECN.

Next, we tested an additional voxel-wise TBSS analysis with average total connectivity beta coefficients as regressors. From that analysis, we found that FA within a small cluster in the medial SLFP was also found to significantly predict total average connectivity of each of the triple-network RSN’s (*p*<0.05, FWE)(*See light blue cluster* **Figure 5**). We also found that FA of a posterior cluster of the right SLFP was positively associated with functional connectivity of the fECN, whereas the medial cluster of the right SLFP was negatively associated with functional connectivity of both the pDMN and coSN (*see* **Figure 5**). We also found that secondary fiber FA in that same right medial SLFT region (i.e. posterior to the DR-associated secondary-fiber cluster described earlier) was positively associated with each of the triple-network RSN’s (R^2^ = 0.35, *F*(3, 32) = 6.65, *p*<0.001; adj. R^2^ = 0.24, *F*(3, 32) = 4.28, *p*=0.001; adj. R^2^ = 0.20, *F*(3, 32) = 3.68, *p*<0.01). That is, localized right SLFP primary and secondary fibers predicted the DR-associated within-network functional connectivity clusters of the triple-network. Since the TRACULA results only provided global measures of FA, averaged across the entire pathway and hence not sensitive to localized clusters, we additionally used the JHU White-Matter atlas to create atlas-labeled ROI’s from the significant TBSS clusters (*p*=0.05 FWE). We could then use these ROI’s to perform a supportive analysis on a non-voxelwise basis. Specifically, this approach would also allow us to explore whether applying atlas definitions to our original TBSS analysis with DR might afford greater specificity into multilayered structural-functional links (See METHODS: TBSS). That analysis further revealed that the DR-associated WM clusters of the CCG, ATR, UF, and Splenium each predicted functional connectivity of the DR-associated triple-network clusters from dual-regression as well (*See* **Appendix, Results: Section A** *for details*). Finally, we tested whether global WM microstructure of the right SLFT, as defined by the tractography measures, might also be driving the multimodal effects (*See* **Figure 6**). The results accordingly showed that average FA of the right SLFT, was positively associated total average intrinsic connectivity of the coSN (adj. R^2^ = 0.13, *F*(3, 29) = 2.62, *p*<0.01) and pDMN (adj. R^2^ = 0.30, *F*(3, 29) = 5.47, *p*=0.001; adj. R^2^ = 0.19, *F*(3, 29) = 3.48, *p*<0.01), but negatively predicted the extrinsic connectivity between the fECN and the left Amygdala (adj. R^2^ = 0.09, *F*(3, 29) = 2.07, *p*<0.05). In sum, both global and local microstructure of right SLFT primary and secondary fibers, along with a variety of other localized WM clusters, predicted within-network functional connectivity patterns of the triple-network associated with DR.

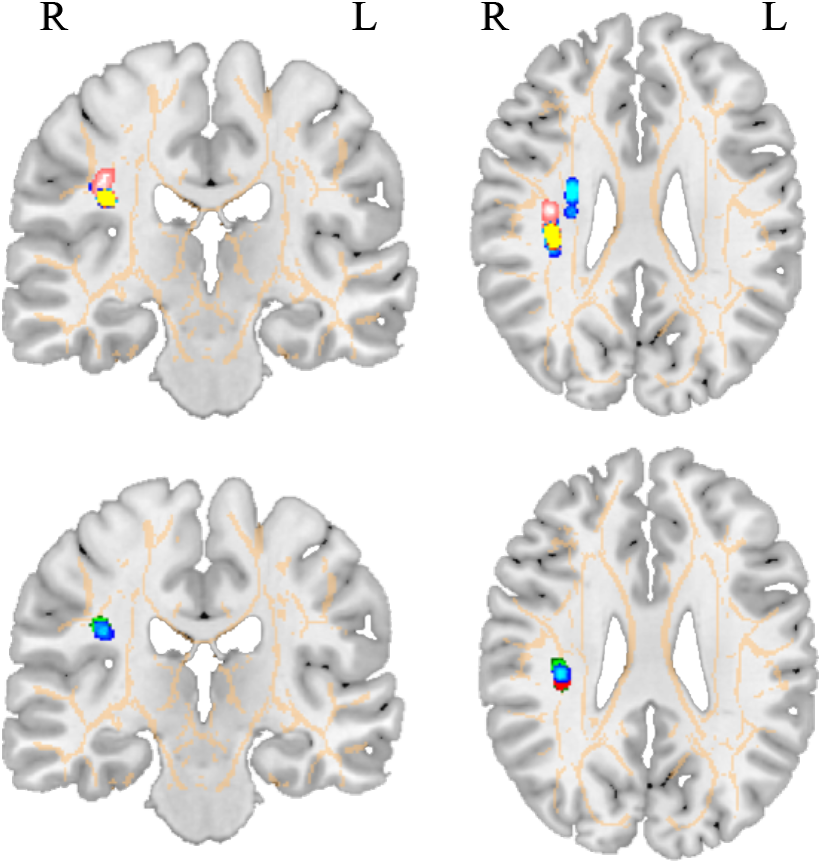

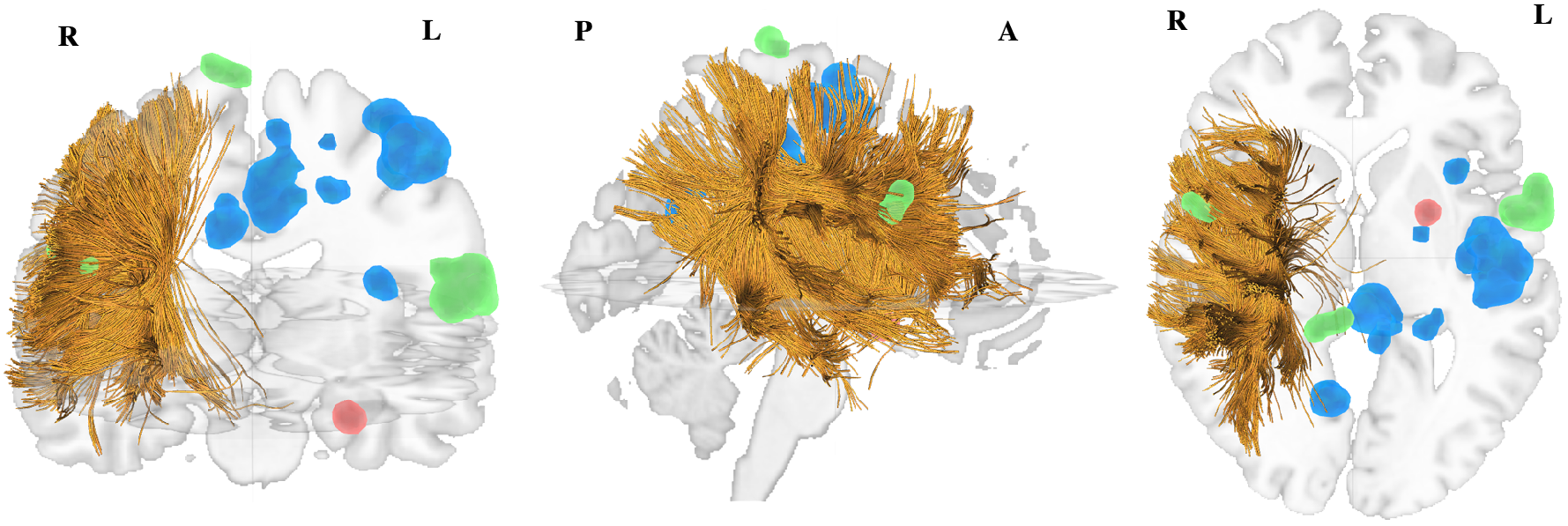

## ‘Between-Network’ Functional Connectivity and Depressive Rumination

### FSLnets

Based on prior evidence for the role of the pDMN, coSN, and fECN in DR, we next used FSLnets to explore whether between-network functional connectivity of each pair combination of triple-network RSN’s was associated with DR (see METHODS: Between-Network Functional Connectivity). That analysis revealed that DR severity was positively associated with an inverse correlation between the fECN and pDMN (*p*<0.05 FWE) (*See* **Figure 7**), and between the coSN and the pDMN (*p*<0.01 FWE). The third ‘between-network’ correlation between fECN and coSN was not significantly associated with DR in the discovery sample.

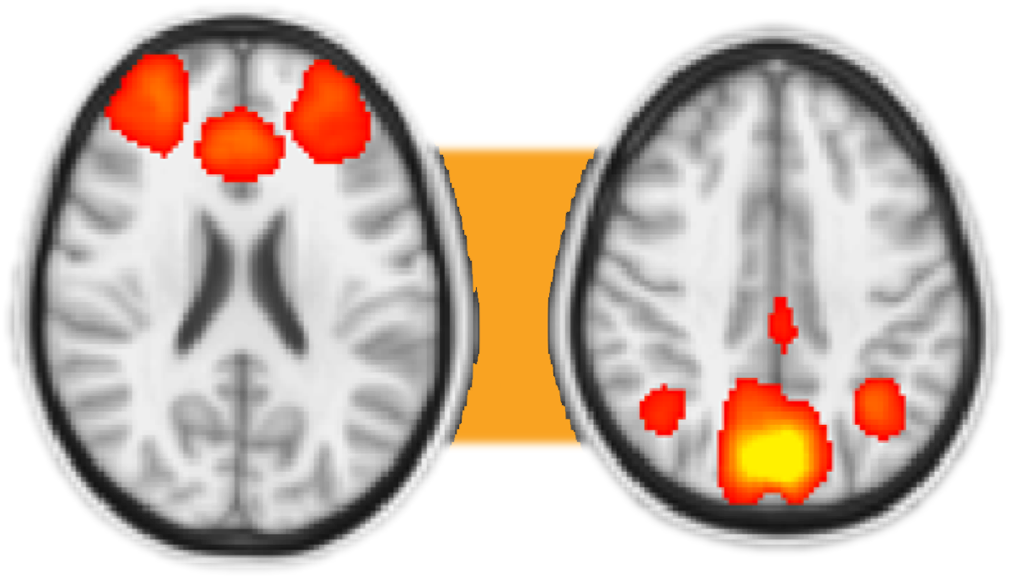

### WM Microstructure and ‘Between-Network’ Functional Connectivity

As in the case of the within-network functional connectivity findings, we also sought to test whether the global and local WM biomarkers of DR predicted the between-network inverse correlations from FSLnets. Accordingly, multimodal analysis revealed that average global FA of the right SLFT, predicted the coSN-pDMN inverse correlation (adj. R^2^ = 0.15, *F*(3, 29) = 2.93, *p*<0.01). Voxel-wise TBSS similarly revealed that the DR-associated coSN-pDMN inverse correlation predicted lower FA of a medial cluster of the right SLFT (adj. R^2^ = 0.17, *F*(3, 32) = 4.17, *p*<0.01) and another along the right posterior Corona Radiata (*p*<0.01 FWE)(*see red/yellow cluster* **Figure 8**). Correspondingly, FA of the left anterior Corona Radiata negatively predicted the pDMN-fECN inverse correlation associated with DR. (*p*<0.05, FWE) (*see blue and red clusters* **Figure 8**). Finally, we again used the ROI’s created from the significant TBSS clusters (*p*=0.05 FWE) to perform a similar non-voxelwise analysis so as to explore whether atlas-defined divisions of WM clusters (i.e. discovered from our original TBSS analysis with DR) might further clarify the nature of their association with between-network functional connectivity. This approach showed that the DR-associated TBSS clusters of the CST, CCG, ATR, UF, and Splenium each predicted the coSN-pDMN inverse correlation, whereas only the left UF predicted the pDMN-fECN inverse correlation (*see* **Appendix, Results: Section B**).

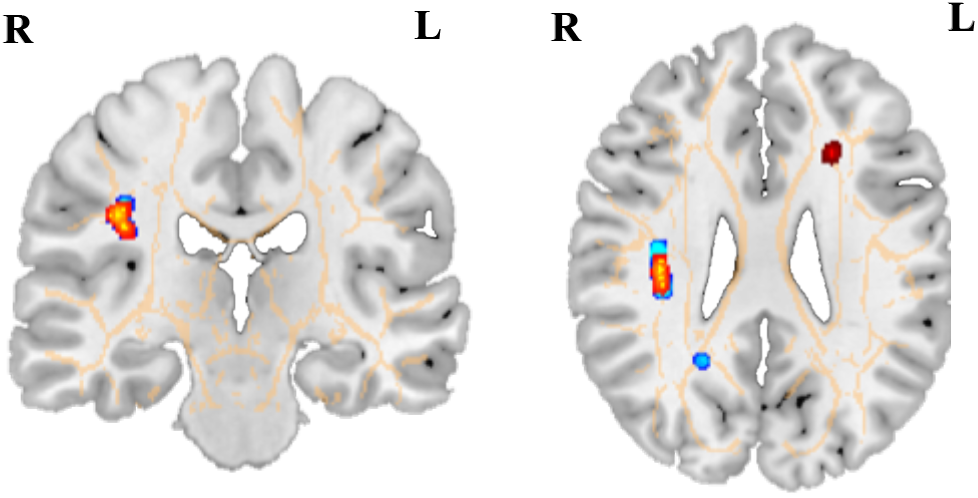

In summary, the initial discovery sample revealed: 1) associations between DR severity and microstructure of the right-lateralized SLFT, along with localized WM clusters scattered throughout the SLFP, CCG, CST, ATR, Splenium, UF; 2) associations between DR severity and the microstructure of secondary fibers in the right superior Corona Radiata; 3) associations between DR severity and both intrinsic/extrinsic within-network functional connectivity of the triple-network; 4) associations between DR severity and between-network functional connectivity of the pDMN with each of the coSN and fECN of the triple-network; 5) associations between localized right medial SLFP primary and secondary-fiber microstructure and each manifestation of triple-network functional connectivity observed to sub-serve DR; 6) an association between right posterior Corona Radiata microstructure and the pDMN - coSN between-network functional connectivity found in (4) to sub-serve DR; 7) associations between left anterior Corona Radiata microstructure and the pDMN – fECN between-network functional connectivity found to sub-serve DR. 8) a variety of less prominent associations between localized WM clusters of the CST, CCG, ATR, UF, and Splenium, along with multiple manifestations of triple-network functional dysconnectivity found in (3) to sub-serve DR. *See* **Figure 9** for a visual summary

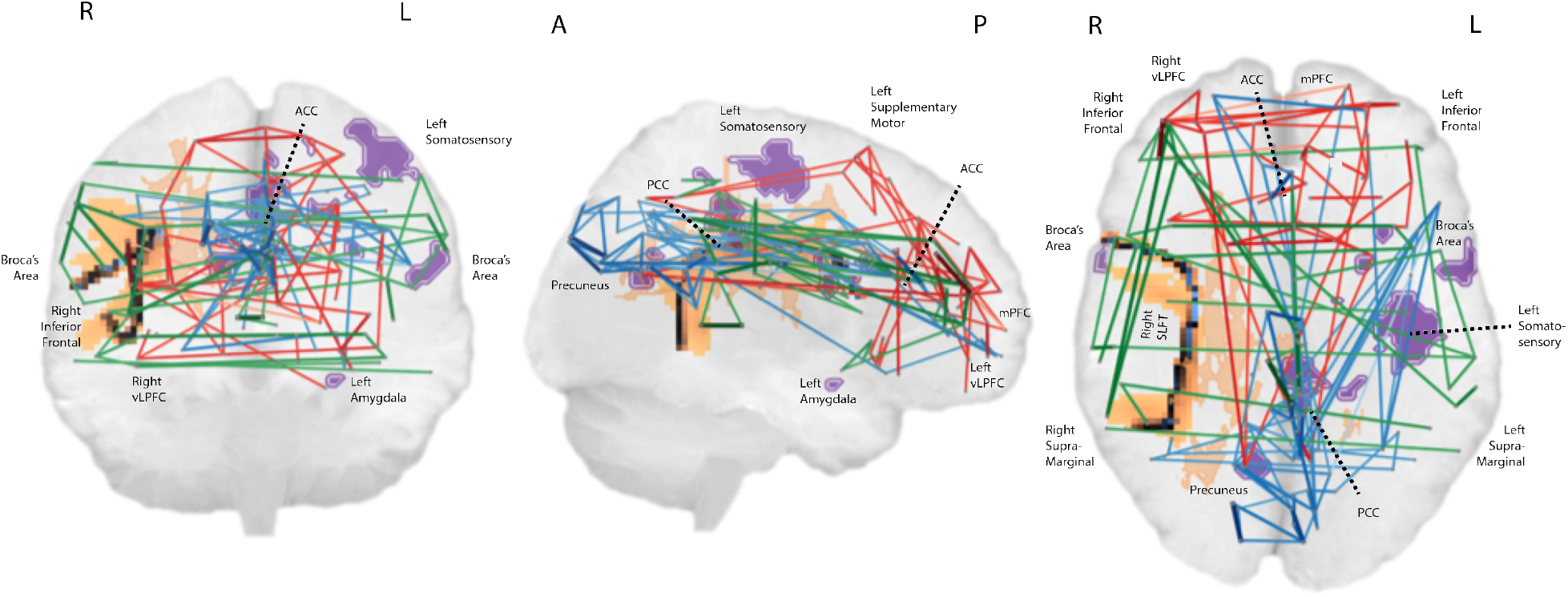

## RESULTS: REPLICATION SAMPLE

The replication sample exhibited similar descriptive statistics to those found in the discovery sample, but with some key differences. Perhaps most notably, for instance, depression severity was considerably milder on average in the replication sample (*M*=10.00, *SD*=6.18, range: 4-31, IQR=5.00). Furthermore, DR severity in the replication sample showed a significant positive skew, which we corrected using a log transformation to yield a normal distribution (Shapiro-Wilk = 0.948, *p* = 0.07). On the other hand, average DR severity overall was similar (*M*=11.00, *SD*=2.74, range: 7-20, IQR=2). Also, age exhibited a slight positive skew in the replication sample (*M*=31.36, *SD* = 5.79, Range: 22-44), as in the discovery sample, along with a similar gender distribution (65% female). *See* **Table 1** for a summary comparing characteristics across samples.

Following the application of a nearly identical analytic methdology to that used with our original sample (*See* METHODS: Replication), we were able to fully replicate 49% of our cumulative findings across all methodologies, modalities, and levels of analysis. Specifically, twenty findings fully replicated, and two findings perhaps ‘partially’ replicated, out of forty retested models from the discovery sample.

**Table 1:**
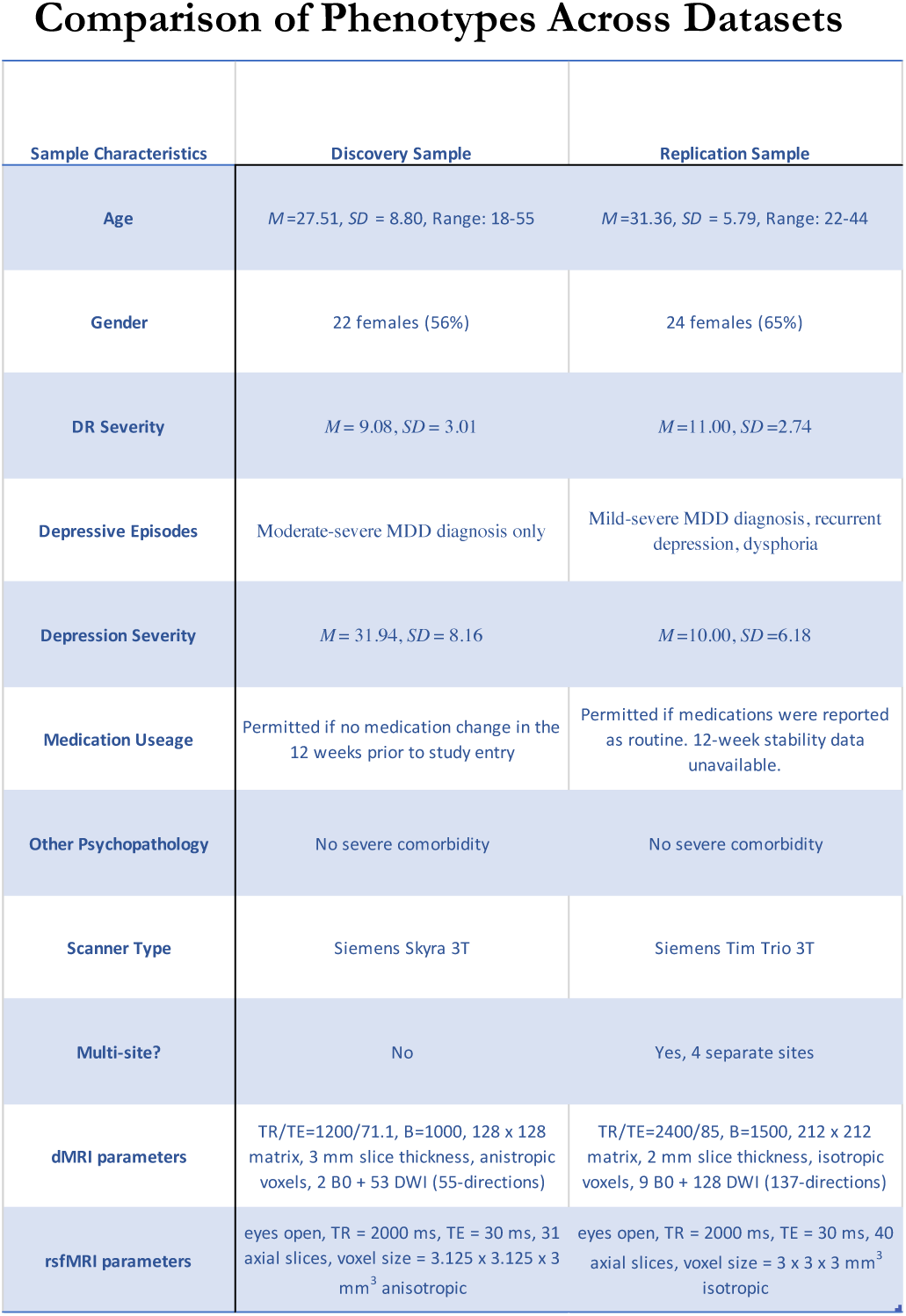
The table above compares key characteristics across samples. In particular, the representation of age, gender, and DR severity was roughly equivalent across samples. On the other hand, the discovery sample included subjects with higher depression severity on average than those included in the replication sample. Although scanner manufacturer and strength was equivalent, the brands differed. Nevertheless, resampling procedures yielded dMRI and rsfMRI data with roughly equivalent core parameters including voxel dimensions, TR, and TE across samples.

| Sample Characteristics | Discovery Sample | Replication Sample |
| --- | --- | --- |
| Age | $M=27.51$, $SD=8.80$, Range: 18-55 | $M=31.36$, $SD=5.79$, Range: 22-44 |
| Gender | 22 females (56%) | 24 females (65%) |
| DR Severity | $M=9.08$, $SD=3.01$ | $M=11.00$, $SD=2.74$ |
| Depressive Episodes | Moderate-severe MDD diagnosis only | Mild-severe MDD diagnosis, recurrent depression, dysphoria |
| Depression Severity | $M=31.94$, $SD=8.16$ | $M=10.00$, $SD=6.18$ |
| Medication Usage | Permitted if no medication change in the 12 weeks prior to study entry | Permitted if medications were reported as routine. 12-week stability data unavailable. |
| Other Psychopathology | No severe comorbidity | No severe comorbidity |
| Scanner Type | Siemens Skyra 3T | Siemens Tim Trio 3T |
| Multi-site? | No | Yes, 4 separate sites |
| dMRI parameters | TR/TE=1200/71.1, B=1000, 128 x 128 matrix, 3 mm slice thickness, anisotropic voxels, 2 B0 + 53 DWI (55-directions) | TR/TE=2400/85, B=1500, 212 x 212 matrix, 2 mm slice thickness, isotropic voxels, 9 B0 + 128 DWI (137-directions) |
| rsfMRI parameters | eyes open, TR = 2000 ms, TE = 30 ms, 31 axial slices, voxel size = 3.125 x 3.125 x 3 mm<sup>3</sup> anisotropic | eyes open, TR = 2000 ms, TE = 30 ms, 40 axial slices, voxel size = 3 x 3 x 3 mm<sup>3</sup> isotropic |

### dMRI Replications (TBSS and Tractography)

The replication yielded a nearly equivalent set of dMRI findings using both TBSS and tractography methods, with a striking degree of similarity in the voxelwise association between WM microstructure and DR severity across samples. Using TBSS, we again found that FA of the right SLFT, SLFP, CCG, CST, Splenium, UF, and ATR exhibited negative correlations with brooding scores (*p*<0.05, FWE)(*See* **Figure 10**). At the *p*=0.05 FWE threshold, 98 voxels, located in the right SLF, CST, and Corpus Callosum (*See top row* **Figure 10**), overlapped in this TBSS analysis across samples. The subsequent ‘crossing-fibers’ TBSS analysis, however, differed considerably across samples (*see* **Table 2** & **Figure 10**). As in the discovery sample, however, the tractography replication demonstrated corresponding negative correlations between DR severity and average FA of localized WM clusters derived from the TBSS ROI-analysis. Additionally, the correlation between DR and global microstructure of the right SLFT was again the only tractography analysis that survived Bonferroni Correction for the six tracts tested (cond. R^2^ = 0.37, *F*(3, 35) = 8.90, p_corrected_<0.05/6 = 0.0083). As in the discovery sample, the effect of hemisphere was again significant with respect to the SLFT (*F*(36,1)=5.43, *p*=0.03), reiterating the right lateralization of this particular global WM biomarker. Whereas in the discovery sample, DR was also positively associated with microstructure of secondary-fibers in a small cluster of the right superior Corona Radiata, in the replication sample, it was negatively associated with microstructure of secondary-fibers of the Corpus Callosum and bilateral CST (p < 0.05 FWE). Although future replications would be needed to confirm, these differences may be largely attributable to the interaction between DR and depression severity across samples (*See* **Appendix, Results: Sections D & E**).

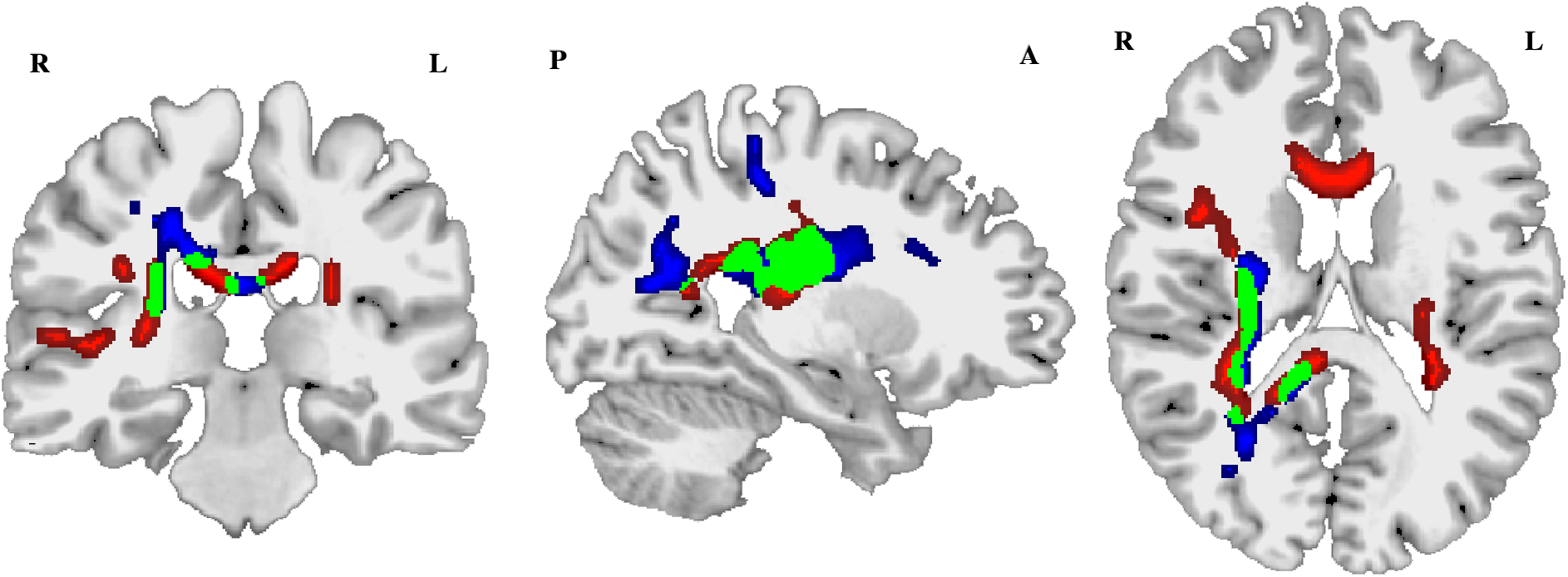

**Table 2:** The above table depicts key findings (*left column*) that directly replicated across the discovery sample (*middle column*) and replication sample (right column). Each of the middle and right columns states the R^2^ value from the respective regression models (adjusted in the discovery sample and conditional on both fixed and random effects in the replication sample), the methodology used to estimate that value (e.g. TBSS ROI-analysis, tractography, dual-regression, FSLnets), and the directionality of the relationship (*neg=negative,pos=positive*). In total, 20 unique findings out of 40 original findings directly replicated across samples.

| Regression Findings (Predictor and Outcome) | Initial Sample | Replication Sample |
| --- | --- | --- |
| Right SLFT Microstructure and DR Severity | TBSS ($p<0.01$ FWE, neg)<br>Tractography ($R^2=0.25$, adj. $R^2=0.18$, neg) | TBSS ($p<0.01$ FWE, neg)<br>Tractography (cond. $R^2=0.37$, neg) |
| Right SLFP Microstructure and DR Severity | TBSS ($p<0.01$ FWE, neg)<br>Tractography ($R^2=0.17$, adj. $R^2=0.05$, neg) | TBSS ($p<0.01$ FWE, neg)<br>Tractography (cond. $R^2=0.23$, neg) |
| CST Microstructure and DR Severity | TBSS ($p<0.01$ FWE, neg)<br>Tractography ($R^2=0.11$, adj. $R^2=0.06$, neg) | TBSS ($p<0.01$ FWE, neg)<br>Tractography (cond. $R^2=0.17$, neg) |
| CCG Microstructure and DR Severity | TBSS ($p<0.05$ FWE, neg) | TBSS ($p<0.01$ FWE, neg) |
| UF Microstructure and DR Severity | TBSS ($p<0.05$ FWE, neg) | TBSS ($p<0.01$ FWE, neg) |
| Splenium Microstructure and DR Severity | TBSS ($p<0.05$ FWE, neg) | TBSS ($p<0.05$ FWE, neg) |
| ATR Microstructure and DR Severity | TBSS ($p<0.05$ FWE, neg) | TBSS ($p<0.05$ FWE, neg) |
| pDMN Within-Network Connectivity and DR Severity | Dual-regression ($p<0.01$, FWE, neg) | Dual-regression ($p<0.001$ unc. ROI-analysis, neg) |
| fECN Within-Network Connectivity and DR Severity | Dual-regression ($p<0.01$, FWE, neg) | Dual-regression ($p<0.001$ unc. ROI-analysis, neg) |
| pDMN-fECN Between-Network Connectivity and DR Severity | FSLnets ($p<0.05$, FWE, neg) | FSLnets ($p<0.01$, FWE, neg) |
| Right SLFT Microstructure and pDMN Within-Network Connectivity | Tractography ($R^2=0.36$, adj. $R^2=0.30$, neg)<br>TBSS ($p<0.05$ FWE, neg) | TBSS ROI-analysis (cond. $R^2=0.36$, neg) |
| Right SLFP Microstructure and pDMN Within-Network Connectivity | Tractography ($R^2=0.26$, adj. $R^2=0.19$, neg)<br>TBSS ($p<0.05$ FWE, neg) | TBSS ROI-analysis (cond. $R^2=0.33$, neg) |
| ATR Microstructure and pDMN Within-Network Connectivity | TBSS ROI-analysis ($R^2=0.33$, adj. $R^2=0.26$, neg) | Tractography (cond. $R^2=0.18$, neg)<br>TBSS ROI-analysis (cond. $R^2=0.28$, neg) |
| CCG Microstructure and pDMN Within-Network Connectivity | TBSS ROI-analysis ($R^2=0.29$, adj. $R^2=0.22$, neg) | Tractography (cond. $R^2=0.12$, neg)<br>TBSS ROI-analysis (cond. $R^2=0.21$, neg) |
| Splenium Microstructure and pDMN Within-Network Connectivity | TBSS ROI-analysis ($R^2=0.21$, adj. $R^2=0.12$, neg) | TBSS ROI-analysis (cond. $R^2=0.23$, neg) |
| Right SLFT Microstructure and coSN Within-Network Connectivity | Tractography ($R^2=0.21$, adj. $R^2=0.13$, neg)<br>TBSS ($p<0.05$ FWE, neg) | Tractography (adj. $R^2=0.50$, neg)<br>TBSS ($p<0.05$ FWE, neg) |
| CCG Microstructure and coSN Within-Network Connectivity | TBSS ROI-analysis ($R^2=0.27$, adj. $R^2=0.20$, neg) | TBSS ROI-analysis (adj. $R^2=0.26$, neg) |
| Splenium Microstructure and coSN Within-Network Connectivity | TBSS ROI-analysis ($R^2=0.17$, adj. $R^2=0.08$, neg) | Tractography (cond. $R^2=0.29$, neg)<br>TBSS ROI-analysis (cond. $R^2=0.55$, neg) |
| Right SLFT Microstructure and fECN Within-Network Connectivity | Tractography ($R^2=0.18$, adj. $R^2=0.09$, neg)<br>TBSS ($p<0.05$ FWE, pos) | Tractography (cond. $R^2=0.31$, pos)<br>TBSS ROI-analysis (cond. $R^2=0.34$, neg) |
| DR-associated WM microstructure and pDMN-fECN Between-Network Connectivity | FSLnets ($p<0.05$ FWE, pos) | FSLnets ($p<0.05$ FWE, pos) |

### rsfMRI Replications (Dual-Regression and FSLnets)

Despite similar microstructural connectivity findings across samples at whole-brain FWE-corrected thresholds *(See green overlapping clusters in* **Figure 10**), the initial within-network functional connectivity findings replicated only when using a small-volume ROI correction (i.e. MNI atlas-defined masks of significant regions from the discovery sample). Nevertheless, the results showed similar within-network functional connectivity patterns of the pDMN and fECN with respect to both anatomical location of intrinsic/extrinsic clusters and directionality (*p*<0.001) (*See correspondence of blue and red clusters in* **Figure 11**). Within the pDMN, lower intrinsic functional connectivity with the right Precuneus replicated (*p*<0.001). And the significant voxels of the pDMN and coSN whose functional connectivity was associated with DR were in close anatomical proximity across samples but not overlapping *(See correspondence of green clusters in* **Figure 11**). In constrast, 9 voxels of the fECN positive connectivity cluster in the left amygdala/parahippocampal gyrus cluster were overlapping. The coSN also neuroanatomically tracked the within-network coSN functional connectivity effects observed discovery sample, but in the reverse direction (*See* **Appendix, Results: Section C** *for a discussion of ‘partial’ replications*).

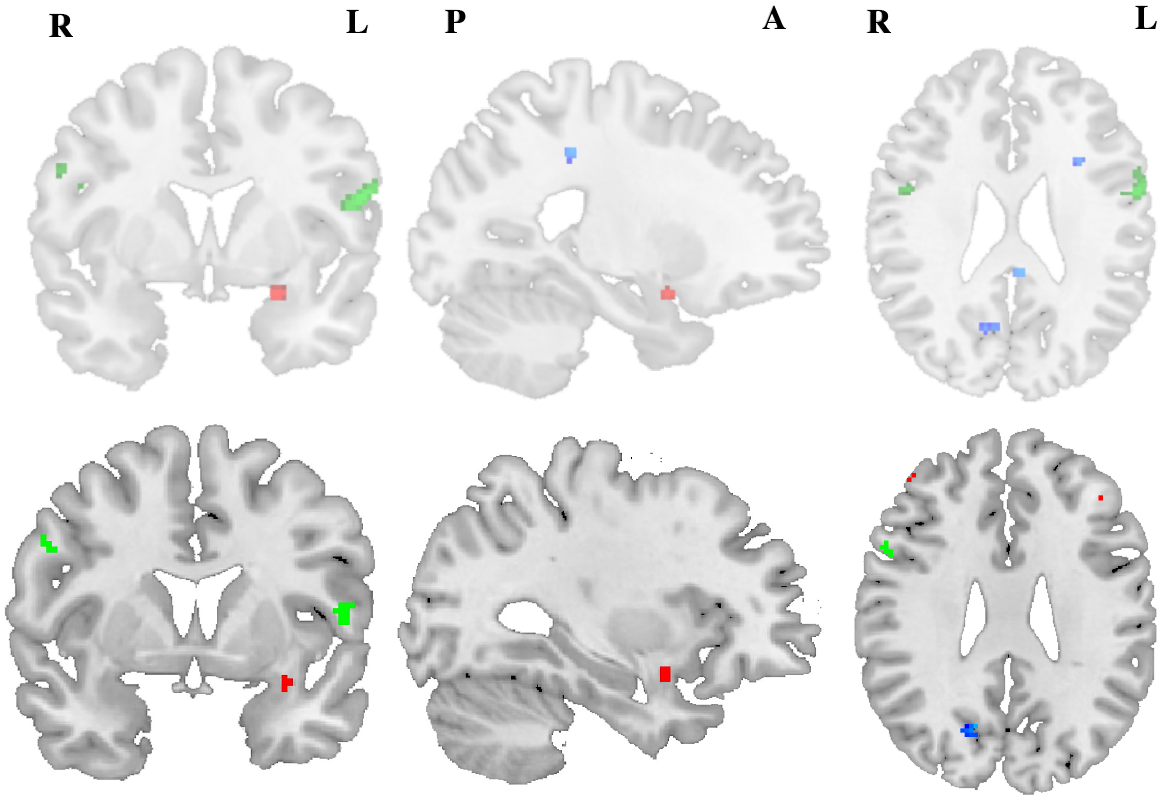

As with the microstructural findings, on the other hand, the between-network functional connectivity findings from FSLnets also replicated with FWE correction. Specifically, the association between DR and the pDMN-fECN inverse correlation fully replicated (*p*<0.05, FWE)(*See* **Figure 12**), whereas the coSN between-network associations with DR diverged across samples (*See* **Appendix, Results: Section C**) in light of the reversed directionality of the coSN’s within-network association with DR.

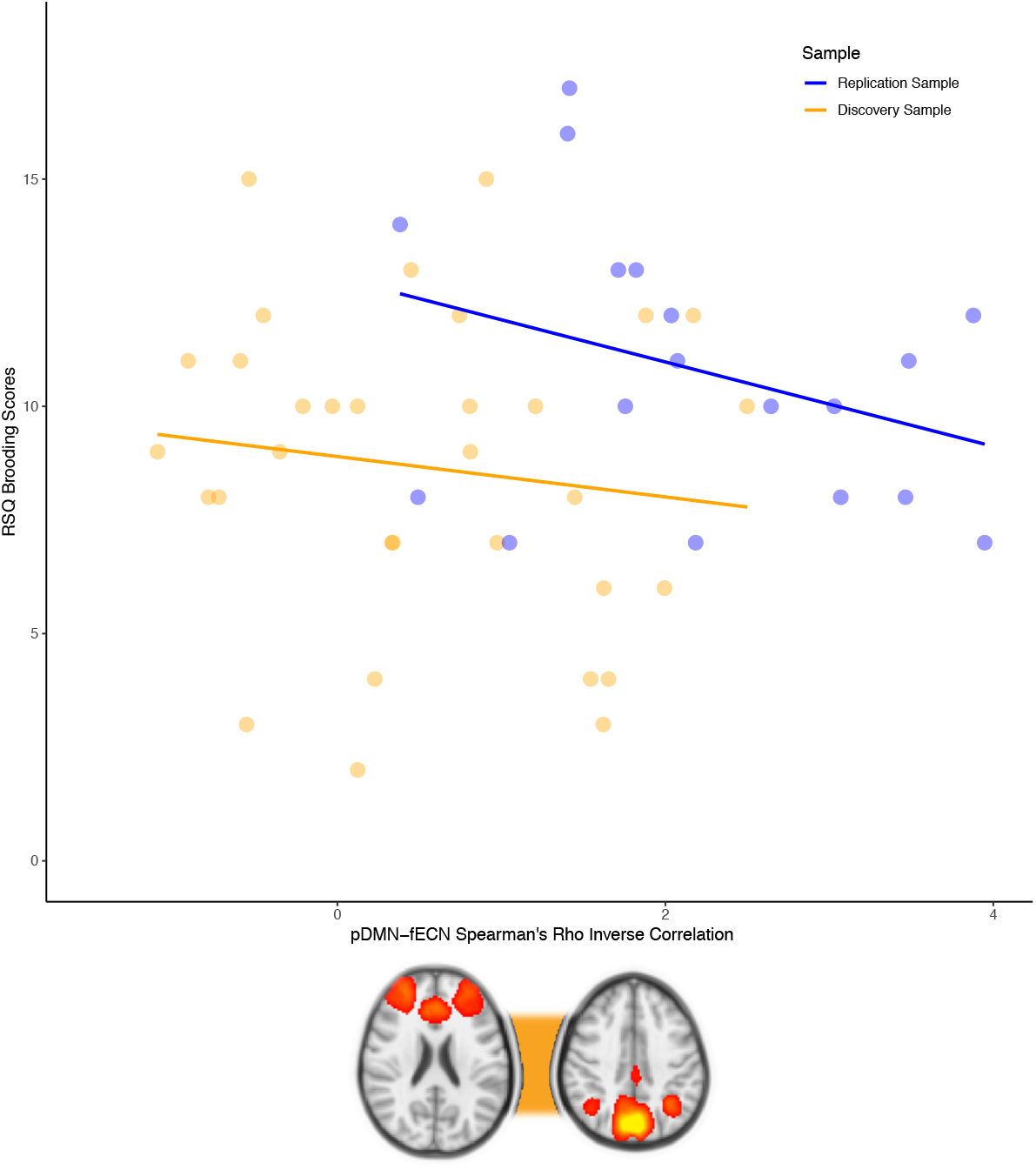

### Multimodal dMRI-rsfMRI Replications

Finally, when exploring multilayer microstructural-functional relationships in the replication sample, we found that global average FA of the right SLFT, along with localized WM clusters of the SLFP, CCG, ATR, and Splenium again predicted both the within-network and between-network functional connectivity biomarkers of DR (*see* **Table 2**). Importantly, the cumulative average FA measure from TBSS predicted pDMN-coSN between-network connectivity in the replication sample (R^2^ = 0.19, *p*<0.05), despite there being no replication of the coSN-pDMN inverse correlation in association with DR. Furterhmore, global microstructure of the right SLFT and localized microstructure of the SLFP, CCG, ATR, and Splenium were again positively associated with the within-network functional connectivity of the pDMN and with corresponding directionality (cond. R^2^ = 0.36, *p*<0.05; cond. R^2^ = 0.28, *p*<0.05; cond. R^2^ = 0.33, *p*<0.05; cond. R^2^ = 0.22, *p*<0.05). The only fECN-related multimodal effect which fully replicated was the association between localized right medial SLFT microstructure and fECN within-network connectivity. In contrast, the relationship between WM microstructure and pDMN-fECN functional connectivity across samples was ultimately non-specific to any single tract and implicated the cumulative DR-associated TBSS clusters, as discovered through a final analysis in FSLnets (*p*<0.05).

We also found that global microstructure of the right SLFT and localized microstructure of the CCG and Splenium were again positively associated with the within-network functional connectivity of the coSN (cond. R^2^ = 0.50, *p*<0.05; cond. R^2^ = 0.26, *p*<0.05; cond. R^2^ = 0.55, *p*<0.05). Despite reversed directionality in DR’s association with coSN within-network functional connectivity, moreover, the directionality of multimodal relationships between WM microstructure and coSN functional connectivity was nevertheless consistent across samples. Notably, secondary-fiber microstructure also appeared to explain the between-network functional connectivity differences across samples. Given that depression severity also differed across samples (*See* **Table 1**), we conducted exploratory analysis to determine whether BDI scores might explain the observed differences in secondary fiber microstructure and coSN functional connectivity across samples (*See* **Appendix, Results: Sections D & E**).

In sum, our results ultimately showed that WM microstructure predicted 24-26% of the variance in DR in both the discovery and replication samples, with the majority of the contribution from global microstructure of the right SLFT. Between-network functional connectivity of the triple-network alone also collectively predicted twice as much variance in DR (50-52%) in both the discovery and replication samples when controlling for WM microstructure. Furthermore, our multimodal results indicated that microstructural connectivity of the SLFP, CCG, ATR, and Splenium in particular largely accounted for the triple-network functional connectivity disturbances associated with DR. Lastly, post-hoc analysis of replication-level False Discovery revealed a 0.1% chance of all multimodal replications being false-positives, and π=0.795 posterior-probability of the replicated large effects, including the ten multilayer microstructural-functional connectivity findings, being true positives (*See* **Appendix, Results: Section F**).

## DISCUSSION

### Overview

The neurobiology of trait Depressive Rumination (DR) is a heterogeneous network phenomenon. Replicated across two independent samples, findings from the present study conceptually converged across multiple dimensions of analysis. With respect to our first hypothesis, we found that compromised resting-state functional connectivity of the brain’s ‘triple-network’ corresponding to self-referential, emotional salience, and attentional engagement processing, forms a functional brain network basis for DR. Likewise, we found that defective global and local frontoparietal White-Matter (WM) pathways in the brain confer microstructural risk factors for DR. Most importantly, however, we observed strong evidence that these microstructural risk factors multimodally perpetuate the trait functional triple-network disorganization that characterizes DR.

Apart from differences in depression severity, discovery and replication samples were matched on key behavioral characteristics: both comprised thirty-nine adults with equivalent measures of DR, similar demographic and inclusion criteria, and corresponding MRI data parameters following resampling procedures (*See* **Appendix, Methods: Sections F & G**). By directly replicating in this way, we created a unique analytic scenario whereby traditional hypothesis testing could be employed to test the generalizability of broader theories beyond individual findings. Principally among the fully-replicated functional connectivity findings, DR severity was negatively associated with within-network intrinsic connectivity of the Precuneal DMN (pDMN) and extrinsic connectivity of the frontal ECN (fECN) with the left Amygdala. DR severity was also negatively associated with a between-network inverse correlation of the pDMN and fECN. On the other hand, DR severity was associated with functional connectivity within the cingulo-opercular SN (coSN) as well as between the coSN and each of the pDMN and fECN; the precise configuration of those relationships differed across samples, however (*See* **Appendix, Results: Sections C & D**).

Microstructurally, DR severity was negatively associated with FA of a temporal section of the right SLFT, a robust finding that replicated across samples. Among the localized WM findings that replicated, we discovered that DR was strongly associated with localized WM clusters of the parietal SLF (SLFP), superior Corona Radiata (SCR), CCG, Corticospinal Tract (CST), Splenium, UF, and Anterior Thalamic Radiation (ATR). DR severity was also positively and negative associated with the microstructure of secondary “crossing” fibers in clusters that were largely localized to key junctures where the aforementioned primary-fiber pathways intersect. These included the Centrum Semiovale, along with the Anterior and Posterior Corona Radiata.

Multimodally, microstructural biomarkers of DR that included global FA of the right SLFT, along with the localized CCG, ATR, and Splenium clusters, predicted the functional connectivity biomarkers of DR, both within and between the pDMN, coSN, and fECN. In contrast, the sample-specific secondary-fiber biomarkers of DR appeared to account for some of the observed variability in coSN functional connectivity across samples, but future studies will be need to confirm this. To our knowledge, this is the first study to show that WM biomarkers of a behavioral measure predict functional connectivity biomarkers of the same behavioral measure.

### WM Microstructure and Depressive Rumination

A growing body of literature on the SLF – the WM biomarker found to most robustly track DR severity– suggests that its microstructure is associated with longer and more severe depressive episodes (Lai & Wu, 2014). Deeper insights might be gleaned, however, from the more fine-grained analysis of the SLF conducted using inter-subject and intra-subject analysis methodologies in tandem (Urgerl et al., 2013). Through both TBSS and tractography, we learned that DR was associated primarily, but not exclusively, with a specific subdivision of the SLF—an inferior portion connecting the middle/superior Temporal Gyrus with ipsilateral prefrontal/cingulo-opercular areas referred to as the SLFT, which most closely corresponds to SLF III – also known as the Arcuate Fasciculus. Neuroanatomically, the SLFT supplies fiber connections between Wernicke’s area for speech production and Broca’s area for language comprehension. The SLFT is most likely responsible for language communication between the parietal and frontal lobes, and specifically for maintaining phonological awareness (Jenkins et al., 2016; Oechslin et al., 2009). Accordingly, research on the microstructural basis of conduction aphasia has shown that SLFT deficits may be related to poor awareness of speech repetition (Bernal & Ardila, 2009). In fact, mammals without vocal repetition ability do not have an SLFT (Schmahmann et al., 2007). As confirmed through tests of “hemisphericity” in both samples, moreover, microstructural deficits of the SLFT associated with DR were also distinctly right-lateralized^i^. Although further research is needed, recent studies have indicated that the right SLF in particular may provide compensatory support for increased demands on language and higher cognitive thought (Glasser & Rilling, 2008; Lindell, 2006). Hence, the DR-associated language deficits of the right SLFT might reflect a lateralization of the WM pathways supporting internal versus external speech. Indeed, cortical areas connected by the right SLFT, such as the right Temporoparietal Junction and right superior Temporal Gyrus, have been implicated in internal speech faculties (Alderson-Day & Fernyhough, 2015; Geva et al., 2011; Nejad et al., 2013). The SLFT deficits observed in the present study may therefore allude to repetition defects of internal speech, and perhaps more broadly, the perseveration of self-referential language of thought itself (Fodor, 1975), as a key maintaining factor for DR (Bernblum & Mor, 2010; Nolen-Hoeksema et al., 1996).

TBSS also revealed a strong association between DR and the microstructure of a variety of localized WM clusters throughout the cortex, which highlights the distributed nature of DR as a microstructural connectivity phenomenon. Additionally, the primary-fiber WM biomarkers were largely consistent across samples, which suggests that these WM deficits in particular are robust risk-factors for DR. In contrast, the secondary-fiber biomarkers revealed by TBSS were largely variable across samples—a discrepancy that may be related to the interaction between depression severity and DR (*See* **Appendix, Results: Sections D & E**), but will require future studies with larger sample sizes to understand fully.

As our study indicated, these DR-associated disturbances in WM follow a subtle mereology. They manifest both locally, whereby they are restricted to distinct clusters of WM, but also globally across the entirety of neuroanatomically distinct fiber bundles such as the SLF. Still, it remains unclear whether the observed WM biomarkers of DR reflect a trait-like vulnerability or develop over the course of extended periods of persistent rumination. In support of the latter possibility, the “perseverative cognition hypothesis” (Brosschot et al., 2006) posits that perseverative cognitive processes like rumination lead to a prolonged stress response, which may further impact myelination patterns in both developing and mature brains (Nugent et al., 2015; Sheikh et al., 2014). To the extent that the perseverative cognition of DR involves repetitive cognitive processing, irregular myelination trajectories might accumulate over time in order to sustain the excess cognitive load inflicted by DR.

### Within-Network Functional Connectivity and Depressive Rumination

The results from our functional connectivity analyses broadly implied that disrupted integration and segregation of self-referential and cognitive control systems are associated with DR. These patterns were firstly observed in the form of disrupted intrinsic and extrinsic functional connectivity within each of the triple-network RSN’s.

#### pDMN

The association between DR and functional connectivity within the pDMN was characterized by two core features: 1) lower intrinsic connectivity with the right Precuneus; and 2) lower extrinsic connectivity with the left Somatosensory Cortex, left inferior Frontal Gyrus, and left Precentral Gyrus. Broadly speaking, diminished Precuneal integration with the DMN may reflect impaired inhibition of internal mentation, particularly given the role of the Precuneus as a core parietal hub for maintaining conscious self-awareness (Klaassens et al., 2017). Conversely, greater segregation from frontal cognitive control modules and somatosensory areas in DR may imply globally dysregulated self-referential processing that is largely detached from present-moment sensory experience (Johnson et al., 2009; Pelletier-Baldelli et al., 2015; Rosenbaum et al., 2017).

#### fECN

Higher DR severity was also associated with greater extrinsic functional connectivity between the fECN and the left Amygdala – a finding that replicated closely across samples and might be explained in a number of ways. One interpretation stems from the premise that DR can be construed as maladaptive coping strategy for avoiding the negative affective states experienced in depression (Watkins & Teasdale, 2001). For example, severe ruminators might repetitively reflect on their own cognitions as a short-term attempt to ward off negative affect, but at the cost of maintaining a dysphoric mood in the long term (Watkins & Brown, 2002). If DR were to accordingly operate *as though* it were adaptive, we might therefore expect that it be accompanied by higher fECN-amygdala functional connectivity as a self-regulatory measure (Cohen et al., 2016). This explanation also aligns with our multimodal findings which indicated that microstructure of the right SLFT may, at least to some extent, reinforce this fECN pattern in a perseverative manner.

#### coSN

DR was also associated with functional connectivity of the coSN, and particularly with clusters corresponding to Broca’s area bilaterally. In light of these clusters being neuroanatomically distinct across samples, the direction of the relationship between DR and the coSN’s patterns of connectivity with these clusters differed across samples. As one explanation for this discrepancy, emerging research offers a compelling case that Broca’s area has two distinct functional subunits: the first supports selective language comprehension and the second is responsible for domain-general cognition (Fedorenko et al., 2012). Whereas the selective language sub-region activates upon the presentation of salient semantic information, the domain-general sub-region activates during more demanding multi-tasking scenarios. Not only can these sub-regions operate more or less independently in response to multiple demands, their precise anatomical boundaries also appear to vary across individuals (Fedorenko et al., 2012; Fedorenko & Varley, 2016). Consequently, the variable functional connectivity between the coSN and each of these versatile sub-regions might indicate multiple “routes” to DR such as excessive negative salience tagging of inner speech and deficient positive salience toward reflective problem-solving. The multimodal findings indicated that global microstructural deficits of the right SLFT actually predicted both of these cases. Likewise, our exploratory tractography revealed that the SLFT fibers terminated in the vicinity of these coSN clusters in the majority of subjects.

### Between-Network Functional Connectivity and Depressive Rumination

Upon discovering spatially-overlapping patterns of aberrant extrinsic functional connectivity among the triple-network RSNs, we further found that DR was associated with patterns of between-network functional connectivity. Specifically, higher DR severity was associated with greater inverse correlation between the pDMN-fECN in both discovery and replication samples, greater inverse correlation between the coSN-pDMN in the discovery sample, and greater direct correlation between the coSN-fECN in the replication sample.

#### pDMN-fECN

Not only did the pDMN-fECN findings replicate those of prior work (Hamilton et al., 2011; Liang et al., 2015), they also replicated across both our initial and replication samples. This finding in particular underscores a consistent pattern of antagonism between self-referential and cognitive control systems as a defining feature of DR. Since the pDMN is functionally coupled with the hippocampus during memory retrieval (Huijbers et al., 2011), greater pDMN-fECN inverse correlation associated with higher DR severity could reflect a failure to regulate the episodic memory refreshing that is believed to occur during self-referential processing (Züst et al., 2015). More broadly, however, greater DMN-ECN coupling likely occurs during task-related autobiographical planning (Spreng et al., 2010) — a pattern that should be antithetical to DR cognitive states. Hence, greater pDMN-fECN inverse correlation associated with DR might conversely implicate the decoupling of cortical midline hubs such as the PCC and ACC, whose synchrony is likely vital to steering self-referential thought in the service of goal-directed cognition (Spreng et al., 2010). When the integration and segregation patterns of these hubs become desynchronized, moreover, negatively-biased internal mentation may become increasingly difficult to regulate (Hamilton et al., 2011).

As with the multimodal relationships discovered when exploring each triple-network RSN separately, tractography also revealed that right SLF microstructure alone predicted the pDMN-fECN inverse correlation in both samples. This finding was somewhat surprising, given that the CCG also supplies direct WM connections between ACC and PCC hubs of both networks. Nevertheless, a recent study found that a disrupted SLF microstructure precluded efficient information transfer between frontoparietal networks in hypertension patients (Li et al., 2015). In fact, the right SLF neuroanatomically supplies lateral frontoparietal WM connections that could impact pDMN-fECN synchronization patterns beyond the influence of more ‘direct’ cortical midline WM pathways.

#### coSN-pDMN and coSN-fECN

Given that the SN is important for switching attention between externally and internally salient stimuli, it is also known to interact with the DMN and ECN across contexts (Menon & Uddin, 2010; Uddin, 2014, 2017). Accordingly, our findings indicated that abnormal between-network relationships with the pDMN and fECN may confer variable risks for DR. As in the case of inconsistent directionality across samples for the coSN within-network connectivity findings, however, the expression of between-network connectivity patterns of coSN with the other triple-network RSN’s also varied across samples. Consequently, this prevented us from forming hard interpretations about its precise role in DR (*See* **Appendix, Results: Sections C, D, & E** *for exploratory analyses*). On the other hand, soft interpretations may be appropriate. Hamilton et al. (2011), for instance, observed a similar pattern of direction reversal with respect to the association between DR severity and between-network activation patterns of the SN as a function of depression severity. Specifically, their state-change analysis showed that relative to healthy controls, depressed participants displayed increased SN activation at the onset of increases in ECN activation relative to the DMN; in contrast, healthy controls exhibited increased SN response at the onset of increases in DMN activation relative to the ECN. Thus, prior evidence suggests that we should expect variable directionality in the relationship between DR severity and the SN’s connectivity profile among the triple-network RSN’s. Although for brevity-sake we only report our analysis of this idea in **Appendix Section E**, we indeed replicated the explanation suggested by Hamilton et al. (2011) that depression severity may moderate the SN’s role among the triple-network RSN’s in route to DR. Nevertheless, we more importantly showed that the multimodal relationships between both coSN-fECN and coSN-pDMN between-network functional connectivity and WM microstructure were consistently localized to the same clusters of right medial SLFT across samples.

### Limitations and Generalizability

One limitation of the present study may be that the neural biomarkers that it uncovered are not purely attributable DR. Given how closely the theoretical constructs of DR and depression severity are intertwined (Roelofs et al., 2006; Treynor et al., 2003), however, there are fundamental theoretical and statistical obstacles to establishing complete specificity of DR biomarkers in any context (Erdur-Bakera & Bugaya, 2010). Nevertheless, the majority of participants included in our replication sample were dysphoric and/or depression-remitted, thereby enabling us to infer that the replicated DR biomarkers are reproducibly observable across multiple depressive disorder subtypes and at varying levels of depression severity.

By validating twenty distinct findings with a replication sample, our study contributes verifiably generalizable DR biomarkers to the larger corpus of depression literature. Still, nearly half of our findings did not replicate—a discrepancy that echoes often-cited concerns regarding reproducibility when working with sub-optimal sample sizes (Eklund et al., 2016; Evans, 2017). Reassuringly, however, those findings that did fully replicate consisted only of large effects that were adequately powered to generalize as such. Importantly, half of these large effects were multimodal and provide fairly decisive evidence that trait DR’s neurobiology is jointly microstructural and functional in nature (Inano et al., 2011; Westlye et al., 2010; Zorlu et al., 2013).

## CONCLUSION

In the present study, we aimed to identify joint microstructural and functional connectivity biomarkers of Depressive Rumination (DR), which we achieved in the form of ten fully-replicated multimodal findings across two independent samples. Converging across inter-subject and intra-subject methods of analysis applied to both datasets, our results firstly showed that microstructural dysconnectivity confers a clear vulnerability for DR. In particular, we found that DR severity is associated with global WM deficits of the right Superior Longitudinal Fasciculus, along with localized deficits in primary-fiber and secondary-fiber WM clusters distributed throughout the cortex. Results from both datasets secondly demonstrated that DR severity is associated with disorganized patterns of within-network and between-network functional connectivity of the ‘triple-network’, comprising the pDMN, coSN, and fECN. Perhaps most strikingly, however, the discovered WM biomarkers multimodally predicted the triple-network functional connectivity biomarkers across both datasets. Collectively, our findings provide strong support for our initial hypotheses and imply that emotionally dysregulated self-referential processing, perpetuated by abnormal attentional and language development, confers multivariate risk factors for trait DR.

Supported by initial discovery and replication, our study embarks on the critical task of delineating a reproducible neurobiological model of DR. Ultimately, we demonstrate that the pathophysiology of trait DR involves a complex neural system characterized by a rich diversity of neural biomarkers. These biomarkers exhibit both global and local ontology, hemisphere-asymmetry and symmetry, intrinsic and extrinsic network properties, as well as multimodality—the collection of which comprises an intricate set of part-whole relations. Future research should therefore caution against reductionistic interpretation when applying exclusively unimodal analyses to DR. Indeed, the quest for its reproducible neurobiology implores that we ‘rethink’ it as an integrated system of brain network mechanisms, whose whole is not merely the sum of its parts.

## ACKNOWLEDGMENTS

The authors would first like to thank the technical staff at the Texas Advanced Computing Center (TACC) for supplying the computational resources that would have otherwise rendered our neuroimaging analyses computationally intractable. We would also like to thank Ryan Hammonds for assistance with data quality control. Additionally, the first author would also like to thank Ari Cohen for her incredible patience and support throughout drafting of the manuscript.

## MATERIALS AND CORRESPONDENCE

All materials and correspondence should be addressed to the first author at

## FOOTNOTES

1 This discrepancy might be explained with respect to individual differences between our English-speaking participants and the Mandarin-speaking participants from the Zeo et al. 2012 study. Specifically, self-referential processing of Mandarin speakers may operate contralaterally to that of English speakers, since more tonal Chinese characters recruit greater right superior Temporal Gyrus activity, whereas more atonal English words do not (Crinion et al., 2009)

